# *De novo* transcriptome sequencing and comparative analysis of midgut tissues of four non-model insects pertaining to Hemiptera, Coleoptera, Diptera and Lepidoptera

**DOI:** 10.1101/127381

**Authors:** Rajesh K. Gazara, Christiane Cardoso, Daniel Bellieny-Rabelo, Clélia Ferreira, Walter R. Terra, Thiago M. Venancio

**Affiliations:** Laboratório de Química e Função de Proteínas e Peptídeos, Centro de Biociências e Biotecnologia, Universidade Estadual do Norte Fluminense Darcy Ribeiro; Av Alberto Lamego 2000, Campos dos Goytacazes, Brazil.; Departamento de Bioquímica, Instituto de Química, Universidade de São Paulo. Av. Prof. Lineu Prestes 748, São Paulo, Brazil.

**Author notes:** Email addresses: RKG, CC, DB-R, CF, WRT *, TMV.

**Keywords:** insects, transcriptomes, whole midgut, anterior midgut, xenobiotics, juvenile hormone

## Abstract

Despite the great morphological diversity of insects, there is a regularity in their digestive functions, which is apparently related to their physiology. In the present work we report the *de novo* midgut transcriptomes of four non-model insects from four distinct orders: *Spodoptera frugiperda* (Lepidoptera), *Musca domestica* (Diptera), *Tenebrio molitor* (Coleoptera) and *Dysdercus peruvianus* (Hemiptera). We employed a computational strategy to merge assemblies obtained with two different algorithms, which substantially increased the quality of the final transcriptomes. Unigenes were annotated and analyzed using the eggNOG database, which allowed us to assign some level of functional and evolutionary information to 79.7% to 93.1% of the transcriptomes. We found interesting transcriptional patterns, such as: i) the intense use of lysozymes in digestive functions of *M. domestica* larvae, which are streamlined and adapted to feed on bacteria; ii) the up-regulation of orthologous UDP-glycosyl transferase and cytochrome P450 genes in the whole midguts different species, supporting the presence of an ancient defense frontline to counter xenobiotics; iii) evidence supporting roles for juvenile hormone binding proteins in the midgut physiology, probably as a way to activate genes that help fight anti-nutritional substances (e.g. protease inhibitors). The results presented here shed light on the digestive and structural properties of the digestive systems of these distantly related species. Furthermore, the produced datasets will also be useful for scientists studying these insects.

## 1. INTRODUCTION

Although insects can be extremely variable in morphology, there is a clear regularity in their digestive functions, which is apparently related to their physiology (Terra, 1990; Terra and Ferreira, 2012). In this regard, the availability of midgut transcriptomes obtained by second-generation sequencing offers a new way to evaluate the differences and similarities of the midgut physiology across divergent groups of insects. With this aim, the midgut transcriptomes of 4 insect species pertaining to 4 different orders were compared here. We provide below a summary of the life styles and available physiological data of the chosen insects, as a basis for discussing results obtained from the transcriptome data.

The yellow meal worm, *Tenebrio molitor* (Coleoptera), is a pest of stored products with worldwide distribution. The *T. molitor* midgut is buffered at pH 5.6 at the two anterior thirds and at pH 7.9 at the posterior midgut (PM) (Terra et al., 1985). Carbohydrate digestion occurs largely at the anterior midgut (AM), whereas proteins are continuously digested by cysteine proteases at the AM and later by serine proteases located at the lumen and surface of PM cells (Terra et al., 1985). A counter current flux of water, powered by secretion of fluid at the PM and absorption at the AM, recovers digestive enzymes from the inside of the peritrophic membrane (i.e. an acellular chitin-protein membrane surrounding the food inside midgut) before excretion (Terra et al., 1985; Cristofoletti et al., 2001).The fall armyworm, *Spodoptera frugiperda* (Lepidoptera), is a pest of several crops, including corn. It is mainly present in subequatorial Africa and Americas (from the south of USA to the south of South America). Digestion in *S. frugiperda* initially occurs in a highly alkaline midgut, mainly by amylases and serine proteases inside the peritrophic membrane (Ferreira et al., 1994a; Ferreira et al., 1994b). Intermediate and final digestive steps occur via the action of enzymes located between the peritrophic membrane and midgut cells, as well as at the surface of midgut cells. A counter current flux of fluid, similar to that described above in *T. molitor*, is also present in *S. frugiperda* (Ferreira et al., 1994a; Ferreira et al., 1994b).

The housefly *Musca domestica* (Diptera) has a worldwide distribution and can carry a variety of pathogens (Ostrolenk and Welch, 1942). The peculiarities displayed by *M. domestica* larvae (as well as by other *Cyclorrhaphous* flies) with regard to their digestive physiology are related to the process of killing and digesting bacteria (Espinoza-Fuentes and Terra, 1987). The midgut of *M. domestica* larvae can be divided in three parts. The middle midgut cells are thought to have an embryonic origin distinct from the AM and PM cells (Poulson and Waterhouse, 1960). The *M. domestica* middle midgut is very acidic, which probably constitutes an adaptation to kill bacteria, the major food source of *M. domestica* larvae. Further, AM amylases deplete the starch content of the meal, making bacteria more susceptible to low pH, digestive lysozymes and cathepsins D of the middle midgut (Espinoza-Fuentes and Terra, 1987). The nutrients from the killed bacteria pass to the PM, where most of the digestion takes place. Therefore, in a sense, the *M. domestica* PM corresponds to the entire midgut of the majority of the other insect larvae (Espinoza-Fuentes and Terra, 1987).

*Dysdercus peruvianus* (cotton-stainer) is a hemipteran that occurs mainly in South America. The midgut of *D. peruvianus* is divided into three sections (V1-V3), which are linked to the hindgut by V4. *D. peruvianus* lacks a chitin-protein peritrophic membrane and have instead a lipoprotein perimicrovillar membrane that ensheathes the midgut cells microvilli as glove fingers (Silva and Terra, 1994). The midgut of *D. peruvianus* is acidic, with a pH of 5.8. V1 is the site of carbohydrate digestion and absorption of water and glucose. V2 and V3 regions are responsible for protein digestion (mainly by cathepsin L, a cysteine protease) and amino acid absorption (Silva and Terra, 1994).

In the present work we report the *de novo* transcriptome sequencing and analysis of midgut tissues of *S. frugiperda* (Lepidoptera), *M. domestica* (Diptera), *T. molitor* (Coleoptera) and *D. peruvianus* (Hemiptera). We used a combined assembly approach to recover longer transcripts, which were annotated with a step-wise pipeline using different databases. This strategy allowed the annotation of a large fraction of the contigs. The sequencing reads were further used to estimate the transcriptional levels of the combined contigs (i.e. unigenes). The integration of functional annotations and transcriptional levels allowed us to find common and divergent aspects in the digestive processes of these four distantly related species, as well as a number of differentially expressed genes and pathways that are critical for midgut physiology and biochemistry.

## 2. MATERIALS AND METHODS

### 2.1 Insects

*S. frugiperda* (Lepidoptera) were laboratory-reared, as previously described (Parra, 1986). *S. frugiperda* larvae were kept in glass vials with the diet at a temperature of 25°C and 14:10 photoperiod (photophase: scotophase). The diet of *S. frugiperda* was composed of kidney bean "carioca" (*Phaseolus vulgaris*), wheat germ, ascorbic acid, yeast, agar and microbial inhibitors. Adults were fed a 10% honey solution. Stock cultures of *D. peruvianus* (Hemiptera) were reared in glass bottles covered with a piece of cloth, under natural photoperiod conditions at a relative humidity of 50-70% at 24 ± 2°C. Insects had access to water and cotton (*Gossypium hirsutum*) seeds that were previously disinfested by freezing. Adults of *D. peruvianus* were used, as this is the stage of more intense feeding. Larvae of *M. domestica* (Diptera) were reared in a mixture of fermented commercial pig food and rice hull (1:2, v/v) (Targa and Peres, 1979). The larvae used in this study were actively feeding individuals at third instar. *T. molitor* (Coleoptera), were bred on wheat bran at 24-26 °C and relative humidity of 70-75%. Fully-grown larvae (each weighing about 0.12 g) of both sexes were used. Only individuals that were feeding actively were selected for sample preparation.

### 2.2 Sample preparation and sequencing

Larvae from *S. frugiperda, M. domestica*, and *T. molitor* and adults of *D. peruvianus* were immobilized on ice and dissected with gloves, sterile forceps and glassware treated with diethyl pyrocarbonate. Pools of individuals were used for RNA extractions, as following: 10 larvae of *S. frugiperda* and *T. molitor*; 20 larvae of *M. domestica* and; 10 females of *D. peruvianus*. Whole midgut (WM) and carcass (larval body without midgut, CAR) were separated. The anterior third of the midgut (i.e. anterior midgut, AM) of *S. frugiperda, T. molitor*, and *M. domestica* and the three last midgut sections (i.e. V2+V3+V4; posterior midgut, PM) of *D. peruvianus* were used. Dissected tissues were kept for a short period in an ethanol-ice bath and stored at -80°C. RNA was extracted using Trizol (Invitrogen). The mRNA was isolated from total RNA with Dynabeads mRNA purification kit (Life technologies). Total RNA and mRNA quality were checked with an Agilent 2100 Bioanalyzer instrument. cDNA libraries (one library per species/tissue) were constructed following Roche recommendations and sequenced on a 454 Genome Sequencer FLX (454 Life Sciences/Roche) at the sequencing facility of the Instituto de Química (University of São Paulo, Brazil). Sequencing reads have been submitted to SRA under the accession number PRJNA388109.

### 2.3 *De novo* transcriptome assembly

The 454 reads were extracted from SFF files using the *sff_extract* program (https://bioinf.comav.upv.es/seq_crumbs/). We used *fastqc* (https://www.bioinformatics.babraham.ac.uk/projects/fastqc) to evaluate sequencing quality and found that the initial and terminal fragments had reduced quality and some technical biases. Therefore, the leading 15 bp of each read were trimmed. The 3´end of each read was also trimmed to keep reads at a maximum length of 500 bp. Reads with less than 100 bp were discarded. These trimming and filtering steps were critical for using a high quality set of reads in the assembly process. We also filtered reads containing some repetitive sequences reported by fastqc. Different 454 libraries of the same species were combined before the assembly step to increase coverage. *De novo* assemblies were performed using Newbler (version 2.5) (Margulies et al., 2005) and MIRA (version 4.0.2) (Chevreux, 2004). It has been demonstrated that significantly improved assemblies can be obtained by merging *de novo* assemblies of 454 reads obtained with different algorithms. In particular, merging of MIRA and Newbler assemblies by CAP3 (Huang and Madan, 1999) has been shown to be the best strategy (Kumar and Blaxter, 2010). In general, each CAP3 contig represents most (if not all) splicing isoforms of a gene; in the present work we use the term “unigenes” when referring to CAP3 contigs. We followed this strategy and assembled our reads independently with Newbler and MIRA; the resulting assemblies were filtered to retain only contigs with at least 100 bp, which were then combined with CAP3 (MIRA and Newbler contigs were treated pseudo-reads by CAP3).

### 2.4 Transcriptome annotation

A stepwise strategy was used to functionally annotate genes as follows: 1) bi-directional BLASTX (Altschul et al., 1997) hit (BBH); 2) Best unidirectional BLASTX hit; steps 1 and 2 were performed using the eggNOG v4.0 arthropod database (Powell *et al*. 2014), with hit coverage of at least 50% of the shortest protein of the pair and e-value < 1e^−3^. Furthermore, contigs were assigned to eggNOG orthologous groups (OGs) using the results from steps 1 and 2; 3) unigenes not annotated in steps 1 and 2 were submitted to a BLASTX search against the Uniref90 database (e-value < 1e^−3^) (Suzek et al., 2007); 4) the remaining unigenes had their longest ORFs extracted with TransDecoder (http://transdecoder.sf.net) and analyzed with HMMER3 (e-value < 1e^−3^) (Eddy, 2011) searches against the PFAM database (Finn et al., 2014). Unigenes were filtered for bacterial contamination by using BLASTX (Altschul et al., 1997) against bacterial proteins from the eggNOG v4.0 database (Powell et al., 2014)(E-value < 1e^−10^). In addition, rRNAs contaminants were removed using BLASTN (E-value<1e^−10^) against all Genbank (Clark et al., 2016) insect rRNA sequences (downloaded on June 26, 2015).

### 2.4 Differential expression analysis and functional enrichment

To estimate transcriptional levels, reads were mapped back to the respective transcriptome assemblies using BWA (Li and Durbin, 2009). Uniquely mapped reads were normalized by “reads per kilobase per million mapped reads” (RPKM). Differentially expressed genes (DEGs) between tissues of the same species (i.e. AM, PM, WM and CAR) were inferred using edgeR (Robinson et al., 2010). Genes with p-value ≤ 10^−3^ and log_2_ (fold change) (log_2_FC) ≥2 or ≤ -2 were considered up-and down-regulated, respectively. Enriched (i.e. over-represented) eggNOG functions in the DEG sets were analyzed using the Fisher Exact Test (p-value ≤ 0.05; Bonferroni-corrected) using R (www.r-project.org).

## 3. RESULTS AND DISCUSSION

### 3.1 *De novo* transcriptome assembly and annotation

A total of 674,972, 772,398, 684,214 and 785,681 reads were sequenced from *D. peruvianus, T. molitor, M. domestica* and *S. frugiperda* libraries, respectively. After quality filtering (see methods for details), 609,407, 677,080, 608,637 and 747,904 filtered reads from *D. peruvianus, T. molitor, M. domestica* and *S. frugiperda*, respectively, were assembled using a combined assembly approach (Table 1; see methods for details). Initial assemblies for all species were obtained by with Newbler 2.5 and MIRA. These *de novo* assemblies were then combined with CAP3, as previously described (Kumar and Blaxter, 2010). Newbler produced 8,093, 12,370, 5,329 and 10,315 contigs of *D. peruvianus, T. molitor, M. domestica* and *S. frugiperda,* respectively (Table 2). MIRA produced in 27,395, 37,703, 20,589, and 28,308 contigs for *D. peruvianus, T. molitor, M. domestica* and *S. frugiperda,* respectively (Table 2). After combining the assemblies for each species using CAP3, we obtained 6,774, 9,583, 4,248 and 7,264 longer and less redundant contigs (i.e. unigenes), for *D. peruvianus, T. molitor, M. domestica* and *S. frugiperda,* respectively (Table 2; Supplementary figure 1). After filtering potential bacterial and rRNA contaminants, we obtained 6,395, 9,010, 4,005 and 6,833 of unigenes for *D. peruvianus, T. molitor, M. domestica* and *S. frugiperda* (Table S1). The final CAP3 transcriptome assemblies can be found on supplementary file 1.

**Table 1:** Sequencing and mapping statistics

| Species | Raw reads | Filtered reads | Total mapped reads | Uniquely mapped reads |
| --- | --- | --- | --- | --- |
| *D. peruvianus* | 674,972 | 609,407 | 531,707 | 445,533 |
| *M. domestica* | 684,214 | 608,637 | 559,459 | 491,359 |
| *S. frugiperda* | 785,681 | 747,904 | 696,223 | 602,167 |
| *T. molitor* | 772,398 | 677,080 | 624,423 | 550,598 |

Note: The mapping statistics were obtained by mapping the filtered reads against the respective transcriptome assembly.

**Table 2:**
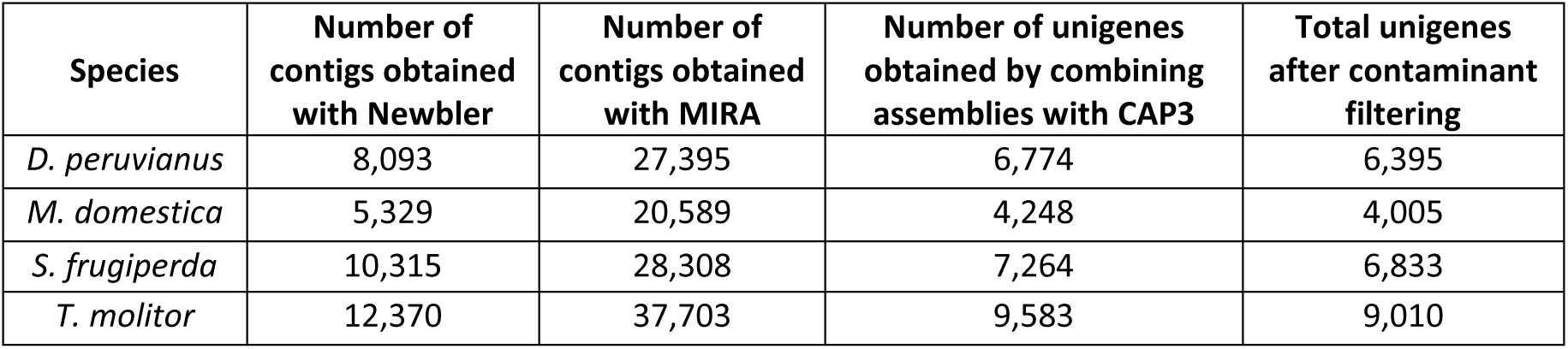
Assembly statistics

EggNOG (Powell et al., 2014) is a database of orthologous groups (OGs) that have general functional annotation codes that are largely derived from the COG database (Tatusov et al., 1997). A stepwise function assignment strategy (see methods for details) allowed us to assign eggNOG function codes for 2,970, 5,764, 2,506 and 3,627 genes from *D. peruvianus, T. molitor, M. domestica* and *S. frugiperda*, respectively (Table 3; Table S2). Through BLASTX against uniref90, 1,809, 2,028, 716 and 1,331 additional unigenes were annotated for *D. peruvianus, T. molitor, M. domestica* and *S. frugiperda,* respectively (Table S2). Finally, the remaining unigenes had their longest ORFs extracted and scanned for conserved PFAM domains, resulting in 317, 595, 299 and 533 additional annotated unigenes from *D. peruvianus, T. molitor, M. domestica* and *S. frugiperda*, respectively (Table S2). In total, we were able to assign some level of functional or evolutionary information to 79.7% to 93.1% of the *de novo* transcriptomes reported here (Table 4).

**Table 3:** Functional classification of eggNOG annotated unigenes

| Classification (one letter eggNOG code) | Number of unigenes (percentages in parentheses) |  |  |  |
| --- | --- | --- | --- | --- |
|  | *D. peruvianus* | *M. domestica* | *S. frugiperda* | *T. molitor* |
| Function unknown(S) | 523(17.61) | 494(19.71) | 894(24.65) | 1255(21.77) |
| Posttranslational modification, protein turnover, chaperones(O) | 398(13.4) | 316(12.61) | 433(11.94) | 690(11.97) |
| Signal transduction mechanisms(T) | 331(11.14) | 205(8.18) | 291(8.02) | 621(10.77) |
| Translation, ribosomal structure and biogenesis(J) | 208(7) | 142(5.67) | 162(4.47) | 225(3.9) |
| Intracellular trafficking, secretion, and vesicular transport(U) | 198(6.67) | 138(5.51) | 154(4.25) | 320(5.55) |
| Carbohydrate transport and metabolism(G) | 169(5.59) | 171(6.82) | 245(6.75) | 426(7.39) |
| Transcription(K) | 160(5.39) | 55(2.19) | 146(4.03) | 331(5.74) |
| Energy production and conversion(C) | 149(5.02) | 63(2.51) | 207(5.71) | 195(3.38) |
| Lipid transport and metabolism(I) | 145(4.88) | 172(6.86) | 175(4.82) | 326(5.66) |
| RNA processing and modification(A) | 130(4.38) | 39(1.56) | 113(3.12) | 207(3.59) |
| Amino acid transport and metabolism(E) | 128(4.31) | 26(1.04) | 241(6.64) | 205(3.56) |
| Cytoskeleton(Z) | 122(4.11) | 106(4.23) | 145(4) | 229(3.97) |
| Secondary metabolites biosynthesis, transport and catabolism(Q) | 93(3.13) | 108(4.31) | 138(3.8) | 197(3.42) |
| Inorganic ion transport and metabolism(P) | 74(2.49) | 65(2.59) | 89(2.45) | 150(2.6) |
| Cell cycle control, cell division, chromosome partitioning(D) | 50(1.68) | 25(1) | 51(1.41) | 116(2.01) |
| Cell wall/membrane/envelope biogenesis(M) | 49(1.65) | 24(0.96) | 34(0.94) | 88(1.53) |
| Nucleotide transport and metabolism(F) | 43(1.45) | 44(1.76) | 63(1.74) | 82(1.42) |
| Coenzyme transport and metabolism(H) | 33(1.11) | 32(1.28) | 35(0.96) | 61(1.06) |
| Replication, recombination and repair(L) | 32(1.08) | 21(0.84) | 54(1.49) | 111(1.93) |
| Chromatin structure and dynamics(B) | 27(0.91) | 10(0.4) | 32(0.88) | 60(1.04) |
| Extracellular structures(W) | 26(0.88) | 16(0.64) | 32(0.88) | 36(0.62) |
| Nuclear structure(Y) | 17(0.57) | 3(0.12) | 8(0.22) | 29(0.5) |
| Defense mechanisms(V) | 16(0.54) | 28(1.12) | 37(1.02) | 54(0.94) |
| Cell motility(N) | 2(0.07) | 2(0.08) | 3(0.08) | 4(0.07) |

Note: A given unigene can be assigned to more than one function code.

### 3.2 Differential gene expression and conservation across species

Next, we further explored the transcriptome datasets reported here to better understand the midgut transcriptional programs. We compared the expression levels of all genes between all possible tissue combinations in each species; PM vs carcass (CAR; whole body except midguts) and PM vs WM, WM vs CAR for *D. peruvianus*; for the remaining species, the following comparisons were performed: AM vs CAR, AM vs WM and WM vs CAR (Figure 1; Table S4). We found a significantly lower number of DEGs between different midgut compartments than when these samples were compared with CAR, which reflect the physiological and anatomical similarities of the midgut compartments. Notably, in *D. peruvianus* there is a relatively small number of induced genes in midgut tissues (Figure 1), probably due to the simplification of the midgut of *D*. *peruvianus* ancestors (insects similar to extant cicads) resulting from their adaption to feed on plant sap, which is generally poor in macromolecules that demand digestion (Terra, 1990). Although other hemipterans regained a macromolecular feeding habit, they did this by deploying lysosomal enzymes instead of the serine proteases that had been previously lost (Houseman et al., 1985). A similar trend has been observed in *M. domestica*, and is in accordance with the streamlined digestive apparatus of the larvae, which is adapted to digest bacterial cells (Levinson, 1960; Lemos and Terra, 1991).

**Figure 1:**
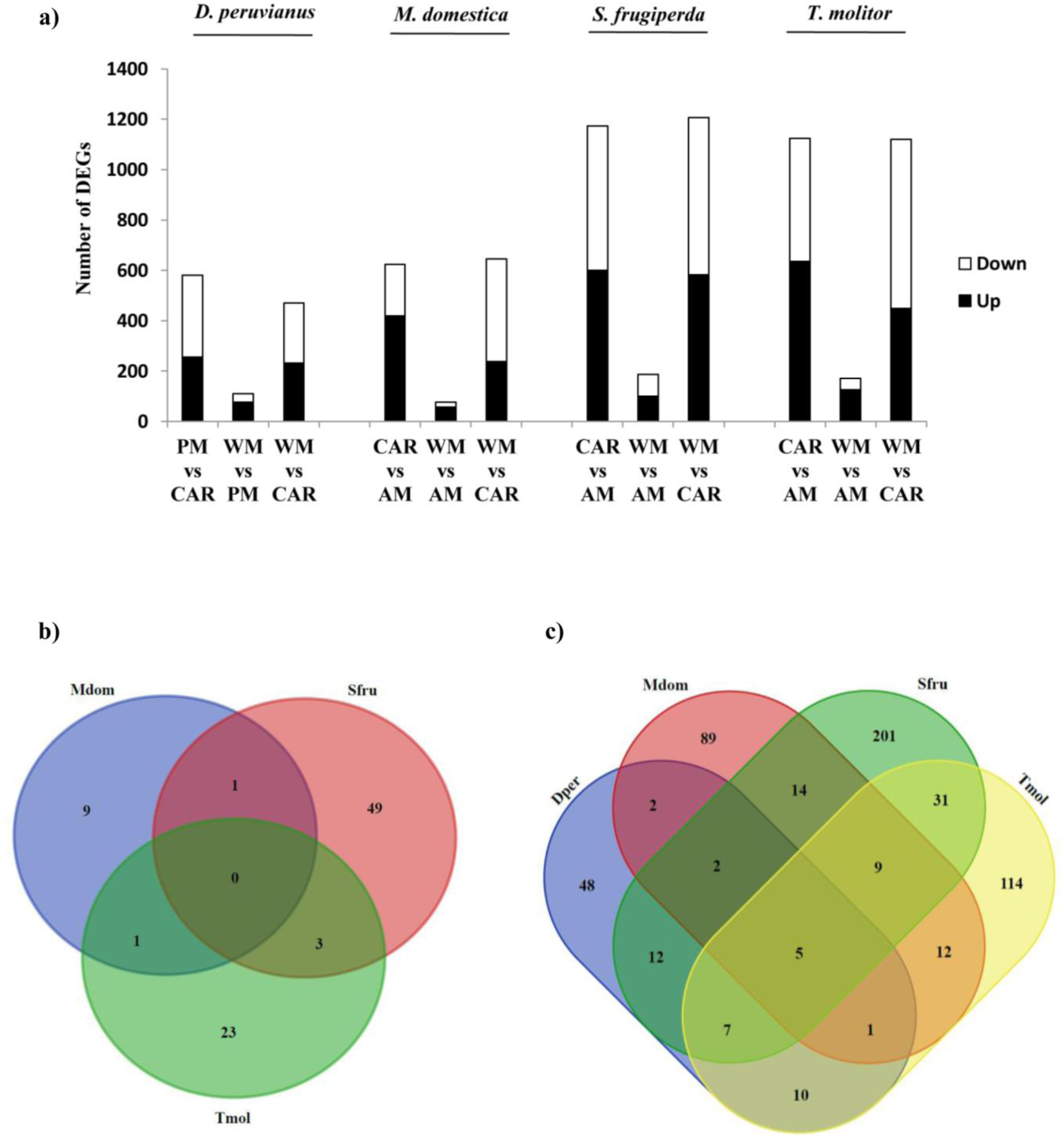
Differentially expressed genes. a) summary of all up (logFC≥2, p-value < =0.001) and down regulated genes (logFC≤2, p-value < =0.001); b) Venn diagram representing eggNOG families with at least one gene from the respective species up-regulated in anterior midgut compared with whole midgut; c) Venn diagram of eggNOG families with at least one gene from the respective species up-regulated in whole midgut compared with carcass. Species abbreviations: Mdom: *M. domestica*, Dper: *D. peruvianus*; Tmol: *T. molitor* and; Sper: *S. frugiperda.*

By aggregating eggNOG family and function annotations to assembled unigenes, we were able to perform functional enrichment analyses to explore the global trends underlying the transcriptomes reported here. In *D. peruvianus*, there is a significant up-regulation of lipases and two endopeptidase families in WM when compared to CAR (Table S4; Table S5). These peptidase gene families encode C1 and D cathepsin family endopeptidases. Interestingly, this latter family is also significantly up-regulated in WM when compared to PM, suggesting that its expression is biased towards the anterior parts of the *D. peruvianus* midgut. It has been hypothesized that up-regulation of cathepsin L (cysteine proteases) and cathepsin D (aspartic proteases) genes are associated with their use as digestive enzymes, replacing lost digestive serine peptidases (Silva and Terra, 1994). Alternatively, it is also possible that those genes were not actually lost, but functionally divergent. This is supported by the down-regulation of three families of serine peptidases (i.e. ENOG4103A6W, ENOG41032G6 and ENOG4103814) in WM (vs CAR). Further, the preferential up-regulation of cathepsin D in the AM is probably an adaptation to remove cathepsin L inhibitors present in the cotton seeds, favoring cathepsin L as the main digestive enzyme in *D. peruvianus* PM (Pimentel et al., 2017). An opposite trend is observed in lipases, which were more expressed in PM than in WM (Table S4; Table S5), supporting a more intense lipid digestion in the posterior parts of the *D. peruvianus* midgut. The up-regulation of lipases could be an adaptation to digest the large oil amounts that are sucked from the cotton seeds (Bewley et al., 2006). Finally, a gene family encoding proteins with a domain of unknown function (DUF3421) is also enriched among genes up-regulated in *D. peruvianus* WM (ENOG410382G). Although the functions of these genes are largely unclear, the two *Drosophila melanogaster* homologs (i.e. CG13321 and CG32633) belonging to this family are highly expressed in embryos/larvae and digestive systems, respectively (modENCODE Anatomy and Development RNA-Seq Atlas, Flybase, accessed on Jan 2017).

In *T. molitor* we found a total of 11 and 2 enriched families among up-regulated DEGs from WM vs CAR and AM vs WM comparisons, respectively (Table S5). These families comprise a number of up-regulated genes involved in digestive and structural functions, such as chitinases, beta-galactosidases, mannosidases, serine-proteases, chitin-binding proteins and peritrophins (Table S4; Table S5). Further, our results suggest that serine protease activity is more constrained to the PM, as these genes are down-regulated in the AM when compared to WM (Table S5). Importantly, the preferential expression of cysteine (e.g. cathepsin L) and serine proteases (e.g. trypsin and chymotrypsin) in the AM and PM of *T. molitor*, respectively, were previously described by direct enzymatic assays (Terra and Cristofoletti, 1996). This strong up-regulation allowed the isolation and characterization many of these digestive enzymes, such as alpha-mannosidases (Moreira et al., 2015), chitinases (Genta et al., 2006), β-1,3-glucanases (Genta et al., 2009), β-glycosidases (Ferreira et al., 2001), β-galactosidases (Ferreira et al., 2003), serine proteases (Cristofoletti et al., 2001) and chymotrypsins (Sato et al., 2008). Furthermore, we found 5 chitin deacetylase-like genes (family ENOG41033EF) that are up-regulated in *T. molitor* WM when compared with CAR, which probably play key roles in determining the peritrophic membrane structure or permeability (Muthukrishnan et al., 2012). Interestingly, a *D. melanogaster* homolog of this gene (CDA9) is also highly expressed in larvae (modENCODE Anatomy and Development RNA-Seq Atlas, Flybase, accessed on Jan 2017).

In *M. domestica*, we also found a general trend for the up-regulation of genes involved in digestive functions in WM and AM. Further, some enzymes encoded by these genes have been characterized, expressed as recombinant proteins and had their 3D structure resolved (Lemos et al., 1993; Cançado et al., 2007). This list comprises, for example, several genes from a lysozyme family (ENOG4103B29) associated with digestion (Table S5). Interestingly, the gene family encoding DUF3421 proteins, up-regulated in *D. peruvianus* WM, is also up-regulated in *M. domestica* WM (vs CAR). In fact, at least one member of this family appears as up-regulated in WM (vs CAR) in all four species analyzed here (Table 5), further emphasizing the importance of ENOG410382G DUF3421 proteins in the insect midguts. The conservation in expression patterns between distant species warrants further biochemical investigations.

**Table 4:** Annotation summary

| Properties | *D. peruvianus* | *M. domestica* | *S. frugiperda* | *T. molitor* |
| --- | --- | --- | --- | --- |
| Total unigenes | 6,395 | 4,005 | 6,833 | 9,010 |
| BBH hit - eggNOG | 2,402 | 1,976 | 2,823 | 4,410 |
| Top BLAST hit - eggNOG | 568 | 530 | 804 | 1,354 |
| Top BLAST hit - uniref90 | 1,809 | 716 | 1,331 | 2,208 |
| Hit with Pfam | 317 | 299 | 533 | 595 |
| Annotated unigenes (total) | 5,096 | 3,521 | 5,491 | 8,387 |
| Annotated unigenes (%) | 79.69 | 87.92 | 80.36 | 93.09 |

**Table 5:** Orthologous groups with at least one member up-regulated in whole midguts when compared to carcasses in all studied insects.

| eggNOG ID | Description |
| --- | --- |
| ENOG410362J | AMOP (MESH genes) |
| ENOG41034JA | glycoside hydrolase family 31 |
| ENOG410382G | Protein of unknown function (DUF3421) |
| ENOG41032RI | Aminopeptidase |
| ENOG41039KJ | Haemolymph juvenile hormone binding protein (JHBP) |

Finally, we found a number of enriched gene families encoding endopeptidases, carbohydrate hydrolases and lipases, among others, as preferentially up-regulated in *S. frugiperda* WM when compared with CAR. Interestingly, three of these families (i.e. ENOG4103AQE, ENOG4103CXY and ENOG41036EN) are biased towards AM regions of the midgut (Table S5). ENOG4103AQE encodes peritrophin-A chitin binding proteins that have crucial roles in larvae of many insects, including the mosquito *A. aegypti* (Venancio et al., 2009). ENOG4103CXY comprises genes encoding alpha/beta hydrolase fold esterases, which are involved in lipid metabolism. Further, this family contains only genes from lepitopterans, including the silkworm *Bombyx mori* and the postman butterfly *Heliconius melpomene*. Finally, we found three genes encoding prostaglandin reductases (ENOG41036EN family) up-regulated in the *S. frugiperda* AM, which might be involved in prostaglandin signaling at this compartment.

Regarding the conservation of up-regulated genes across different species, we also found some interesting trends, such as: 1) the absence of shared families with up-regulated members in AM (vs WM) of *M. domestica, S. frugiperda* and *T. molitor* (the transcriptome of *D. peruvianus* AM has not been sequenced), which is probably due to different factors, such as the diversification of this tissue across species (e.g. *M. domestica*) and the similarity between the WM and AM transcriptional programs. Importantly, we were able to detect DEGs in AM (vs WM), suggesting that this result is not merely due to inadequate tissue dissection or absence of differential expression throughout the midgut; 2) there are 5 gene families with at least one up-regulated member in the WM of each species (Table 5). These families encode some interesting functions, such as the haemolymph juvenile hormone (JH) binding protein (JHBP, family ENOG41039KJ, discussed in the next section). Other interesting family is ENOG410362J, which encodes orthologs of the *Drosophila* MESH gene (FBgn0051004). In *D. melanogaster*, MESH is highly expressed after 12hr of embryo development and is essential for the development of 1^st^ instar larvae, probably due to its crucial roles in septate junction formation (Izumi et al., 2012). Thus, the up-regulation of MESH genes in such distantly related species reveal the importance of this ancient structural component in larval midgut development; 3) the up-regulation of orthologous families involved in the catabolism of xenobiotics across species. One notable example is a family of UDP-glucoronosyl and UDP-glucosyl transferases [ENOG41033B3; PFAM PF00201 (UDPGT)], with at least one up-regulated gene in the WM of *D. peruvianus, S. frugiperda* and *T. molitor*. This domain is present in a relatively big family of enzymes involved in cuticle formation, pigmentation, olfaction and detoxification of xenobiotics. This latter function is of particular importance to the midguts, which are the primary sites of digestion and are in constant contact with deleterious substances (Heidel-Fischer and Vogel, 2015). Interestingly, in *D. melanogaster*, there are 10 UDPGT genes (8 from the ENOG41033B3 family) in a region of ∼51.5Kb at chromosome 3R (from 11,126,480 to 11,178,000), strongly supporting the expansion of this family via tandem (proximal) gene duplication. Furthermore, a family of cytochrome P450 genes (ENOG41035SH), known to be involved in xenobiotic detoxification (Heidel-Fischer and Vogel, 2015), is also up-regulated in the WM of *D. peruvianus* and *S. frugiperda*, unveiling an additional component of what seems to be an ancient defense program that is activated in the midgut of distantly related species.

### 3.3 Conservation of the top highly expressed genes

Next, we analyzed the conservation patterns of the most highly expressed genes in each tissue in the 4 species considered here. In spite of the evolutionary distance between these species, we found in WM and CAR, respectively, a total of 35 and 36 gene families with at least one conserved member among the top 10% most highly expressed genes in each species (Figure 2 and Tables 6 and S6). Among those 35 families with highly expressed genes in WM, 22 also have highly expressed members in CAR and are probably housekeeping genes. The remaining 13 families probably play more specific roles in the digestive tract. These families comprise some interesting functions, such as peptidoglycan recognition proteins (ENOG41032AN), which is involved in innate immunity of the moth *Trichoplusia ni* (Kang et al., 1998). CG9380, a *D. melanogaster* homolog from this family, displays high expression levels in larval and adult digestive systems (modENCODE Anatomy and Development RNA-Seq Atlas, Flybase, accessed on Jan 2017). Interestingly, no genes from this family are among the top 10% expressed genes in AM, indicating that their functions could be more relevant in more posterior midgut sections. Another interesting gene family showing biased transcription in the midgut of several species is ENOG41039KJ, encoding the abovementioned JHBPs, which protect the JH molecules from hydrolysis in the hemolymph. Furthermore, JH and JHBP can help to counter the effects of anti-nutritional substances (e.g. protease inhibitors) via up-regulation of midgut proteases, as shown in the migratory locust *Locusta migratoria* (Spit et al., 2016). JH/JHPB has also been demonstrated to induce the expression of proteases in other insects, such as *Helicoverpa armigera* (Sui et al., 2009), *A. aegypti* (Sui et al., 2009) and *Hodotermopsis sjostedti* (Cornette et al., 2013). Finally, we have also found a family of serine carboxypeptidases (i.e. ENOG410351F) in this dataset. This is interesting because most digestive carboxypeptidases in these insects (except for *D. peruvianus*, which does not have digestive metallocarboxypeptidases) are metallocarboxypeptidases, whereas the serine carboxypeptidases are mainly lysosomal (Ferreira et al., 2015). This result suggests that the recruitment of lysosomal carboxypeptidases to digestive processes predates the divergence of these insect groups. However, more analyses are warranted to rule out the possibility of independent events after the radiation of these orders.

**Figure 2:**
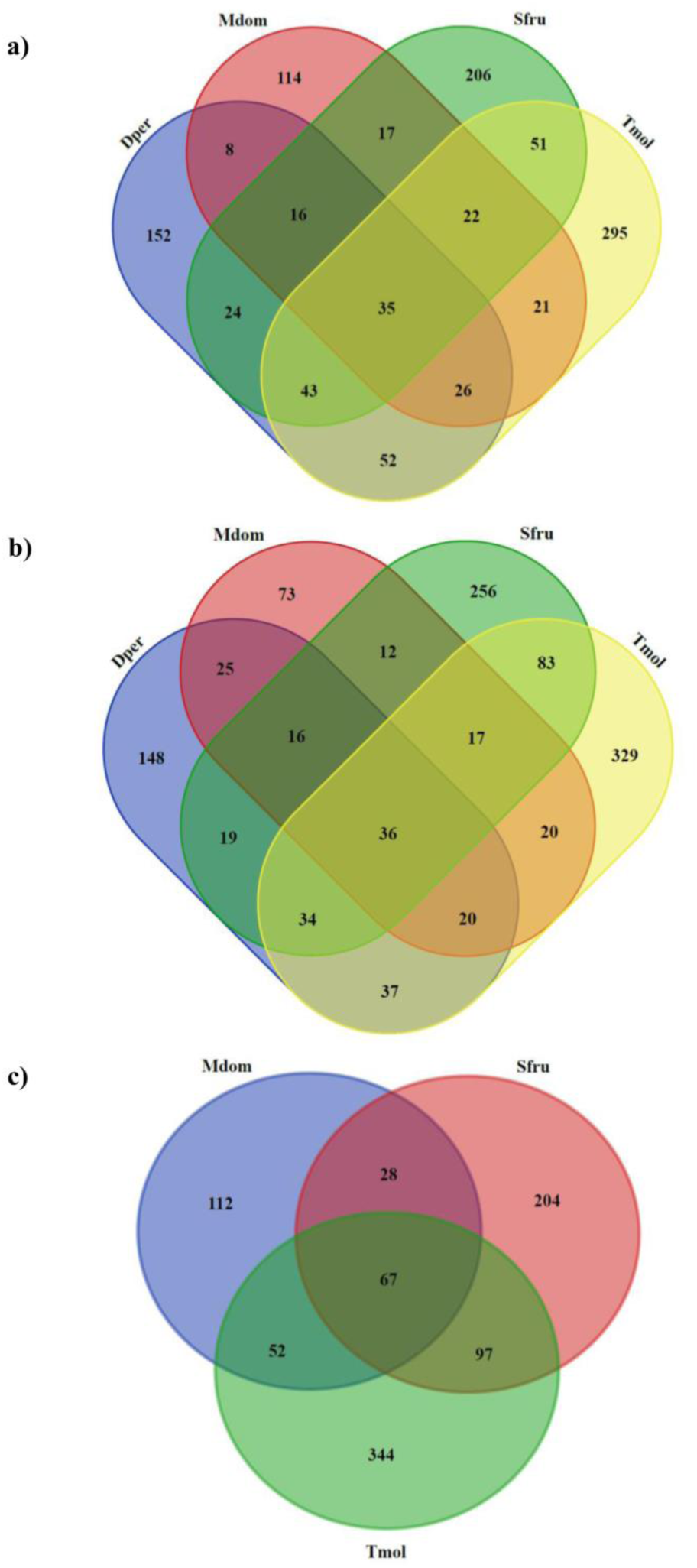
Venn diagram representing eggNOG families with at least one gene among the 10% most highly expressed genes in the respective species in: a) whole midgut; b) carcass; c) anterior midgut. Species abbreviations: Mdom: *M. domestica*, Dper: *D. peruvianus*; Tmol: *T. molitor* and; Sper: *S. frugiperda.*

**Table 6:** Orthologous groups with at least one member among the top 10% most highly expressed unigenes in whole midgut.

| eggNOG ID | Description | Dper | Mdom | Sfru | Tmol |
| --- | --- | --- | --- | --- | --- |
| ENOG41031YA | ATPase subunit B | Y | Y | Y | N |
| ENOG4103206 | ribosomal protein L19 | Y | Y | Y | Y |
| ENOG4103269 | ribosomal protein S9 | Y | Y | Y | Y |
| ENOG41032AN | NA; possibly involved in peptidoglycan recognition | N | N | N | N |
| ENOG41032F0 | ribosomal protein P0 | Y | Y | Y | Y |
| ENOG41032JU | vacuolar ATP synthase subunit H | N | N | Y | N |
| ENOG41032Q3 | Tubulin is the major constituent of microtubules. | Y | Y | Y | Y |
| ENOG41032RI | Aminopeptidase | N | N | N | N |
| ENOG4103369 | activated protein kinase c | Y | Y | Y | Y |
| ENOG410336G | citrate synthase | Y | Y | Y | N |
| ENOG410338D | ribosomal protein L5 | Y | Y | Y | Y |
| ENOG41033B8 | Mitochondrial membrane ATP synthase | Y | Y | N | Y |
| ENOG41033DB | Protein disulfide isomerase | Y | Y | Y | Y |
| ENOG41033N4 | Ribosomal protein L15 | Y | Y | Y | Y |
| ENOG41033P7 | ribosomal protein | Y | Y | Y | Y |
| ENOG41034RS | Sterol carrier protein | N | N | Y | Y |
| ENOG41034Y5 | Involved in calcium binding and microtubule stabilization (By similarity) | Y | Y | Y | Y |
| ENOG410351T | 40S ribosomal protein S8 | Y | Y | Y | Y |
| ENOG4103520 | Thiolase, C-terminal domain | N | N | N | N |
| ENOG4103523 | polyadenylate-binding protein | Y | Y | Y | Y |
| ENOG410356S | Cathepsin | Y | N | Y | Y |
| ENOG4103598 | 40S ribosomal protein | Y | Y | Y | Y |
| ENOG410359F | This protein promotes the GTP-dependent binding of aminoacyl-tRNA to the A-site of ribosomes during protein biosynthesis (By similarity) | Y | Y | Y | Y |
| ENOG41035BZ | Produces ATP from ADP in the presence of a proton gradient across the membrane (By similarity) | Y | Y | Y | Y |
| ENOG41035IX | Tubulin is the major constituent of microtubules. | Y | Y | Y | Y |
| ENOG41035KN | Required for the assembly and or stability of the 40S ribosomal subunit. | Y | Y | Y | Y |
| ENOG4103648 | Arginine kinase | Y | Y | Y | Y |
| ENOG41036EA | ribosomal protein S3 | Y | Y | Y | Y |
| ENOG41036JR | Ribosomal protein S16 | Y | Y | Y | Y |
| ENOG41036K1 | Ribosomal protein S11 | Y | Y | Y | Y |
| ENOG410373X | Tetraspanin family | N | N | N | N |
| ENOG410375W | SERine Proteinase Inhibitors | N | Y | Y | Y |
| ENOG41037D0 | cytochrome C oxidase | N | N | Y | Y |
| ENOG410382G | Protein of unknown function (DUF3421) | N | N | N | Y |
| ENOG41039KJ | Haemolymph juvenile hormone binding protein (JHBP) | N | N | N | N |

The last four columns indicate whether the gene family is also highly expressed in carcass, which is a strong feature of housekeeping genes. Species abbreviations: Mdom: *M. domestica*, Dper: *D. peruvianus*; Tmol: *T. molitor* and; Sper: *S. frugiperda*.

There are 67 gene families with members among the 10% most expressed genes in the AM of *M. domestica, S. frugiperda* and *T. molitor* (AM of *D. peruvianus* was not studied) (Figure 2 and Table 7). Although most of these families encode genes that are also highly expressed in CAR, some of them are obviously related to digestive functions, such as: alpha-amylase, aminopeptidase, alkaline phosphatase, serine-type carboxypeptidase, cathepsin, glucosylceramidase, serine endopeptidase, and serpins (Table 7). This is probably due to the fact that carcasses include the Malpighian tubules, which are known to harbor several enzymes and transporters that are also found in the midgut (Beyenbach et al., 2010). A similar phenomenon occurs with mammalian kidneys and intestines (Boron and Boulpaep, 2005). Further, among the 10% most expressed genes, *T. molitor* and *S. frugiperda* share more OGs with one another than the latter with *M. domestica* (Figure 2). Although unexpected given the phylogenetic relationships of these species, this result can be explained by the adaptation of *M. domestica* to digest bacteria (Levinson, 1960; Lemos and Terra, 1991), which largely remodeled its digestive enzymatic apparatus. Further, most of these 67 gene families encode enzymes involved in primary metabolism and ribosomal proteins. However, important regulatory gene families are also present. A notable example is the ENOG41038D7 family of bZIP transcription factors, which includes the *D. melanogaster* ATF4 (CRC, *cryptocephal*). CRC is involved in critical developmental processes, such as larval molting and pupariation (Hewes et al., 2000), probably in a pathway involving the transcriptional regulation of the Ecdysis triggering hormone (ETH) in endocrine cells (Gauthier and Hewes, 2006). Interestingly, ATF4/CRC is also involved in endoplasmic reticulum stress response through the regulation of a metabolic shift involving the up-regulation of genes encoding glycolytic enzymes (Lee et al., 2015), several of which being present in the set of 67 gene families mentioned above (i.e. ENOG41032XF: Triose-phosphate isomerase; ENOG41035ZK: Glyceraldehyde-3-phosphate dehydrogenase). Therefore, important parts of this system are present in high levels in the AM of different species, suggesting that the regulatory mechanism outlined above play important roles in the development of this tissue across different orders.

**Table 7:** Common OGs with at least one member in the top 10% most highly expressed genes in anterior midgut.

| eggNOG_cID | Description | Mdom |  | Sfru |  | Tmol |  |
| --- | --- | --- | --- | --- | --- | --- | --- |
|  |  | WM | CAR | WM | CAR | WM | CAR |
| ENOG41031YA | ATPase subunit B | Y | Y | Y | Y | Y | N |
| ENOG4103206 | ribosomal protein L19 | Y | Y | Y | Y | Y | Y |
| ENOG4103269 | ribosomal protein S9 | Y | Y | Y | Y | Y | Y |
| ENOG41032DQ | Calreticulin | N | N | Y | Y | Y | Y |
| ENOG41032F0 | ribosomal protein P0 | Y | Y | Y | Y | Y | Y |
| ENOG41032FD | ribosomal protein | Y | Y | Y | Y | Y | Y |
| ENOG41032FU | K05692 actin beta gamma 1 | Y | Y | Y | Y | Y | Y |
| ENOG41032Q3 | Tubulin is the major constituent of microtubules. | Y | Y | Y | Y | Y | Y |
| ENOG41032QK | alpha-amylase | Y | Y | Y | N | Y | Y |
| ENOG41032QY | Peroxiredoxin (EC 1.11.1.15) | N | N | N | N | N | Y |
| ENOG41032RI | Aminopeptidase | Y | N | Y | N | Y | N |
| ENOG41032XF | Triose-phosphate isomerase | Y | N | Y | Y | Y | Y |
| ENOG41032ZC | Angiotensin-converting enzyme | Y | Y | Y | N | Y | Y |
| ENOG4103369 | activated protein kinase c | Y | Y | Y | Y | Y | Y |
| ENOG410336G | citrate synthase | Y | Y | Y | Y | Y | N |
| ENOG4103370 | Occurs in almost all aerobically respiring organisms and serves to protect cells from the toxic effects of hydrogen peroxide (By similarity) | Y | Y | Y | Y | Y | Y |
| ENOG410338D | ribosomal protein L5 | Y | Y | Y | Y | Y | Y |
| ENOG41033AG | Phosphopyruvatehydratase(EC 4.2.1.11) | N | Y | Y | Y | N | Y |
| ENOG41033B3 | Glucuronosyltransferase (EC 2.4.1.17) | N | N | Y | N | Y | N |
| ENOG41033B8 | Mitochondrial membrane ATP synthase | Y | Y | Y | N | Y | Y |
| ENOG41033DB | Protein disulfide isomerase | Y | Y | Y | Y | Y | Y |
| ENOG41033DK | oxidoreductase activity | Y | N | Y | N | Y | N |
| ENOG41033E4 | Ribosomal protein L12 | Y | Y | Y | Y | Y | Y |
| ENOG41033N4 | Ribosomal protein L15 | Y | Y | Y | Y | Y | Y |
| ENOG41033P7 | ribosomal protein | Y | Y | Y | Y | Y | Y |
| ENOG41033ZE | Actins | N | Y | Y | Y | Y | Y |
| ENOG410340Q | Malate dehydrogenase | Y | Y | Y | Y | Y | Y |
| ENOG4103418 | Insulinase (Peptidase family M16) | N | N | Y | Y | Y | Y |
| ENOG410347N | Aldehyde dehydrogenase | Y | N | Y | Y | N | Y |
| ENOG41034BW | Phosphate carrier protein | N | Y | Y | Y | Y | Y |
| ENOG41034I7 | Produces ATP from ADP in the presence of a proton gradient across the membrane (By similarity) | N | Y | Y | Y | Y | Y |
| ENOG41034JA | Glycosyl hydrolase family 31 | N | N | Y | N | N | N |
| ENOG41034MU | 5'-nucleotidase, C-terminal domain | Y | N | Y | N | Y | N |
| ENOG41034RS | Sterol carrier protein | Y | N | Y | Y | Y | Y |
| ENOG41034UM | Enoyl-CoA hydratase | N | Y | Y | Y | Y | Y |
| ENOG41034Y5 | Involved in calcium binding and microtubule stabilization (By similarity) | Y | Y | Y | Y | Y | Y |
| ENOG41034Z5 | fructose-bisphosphatealdolase | N | Y | Y | Y | Y | Y |
| ENOG410351F | serine-type carboxypeptidase activity | Y | N | Y | N | Y | N |
| ENOG410351T | 40S ribosomal protein S8 | Y | Y | Y | Y | Y | Y |
| ENOG410351Z | peroxisomal multifunctional enzyme type | Y | N | Y | Y | Y | N |
| ENOG4103520 | Thiolase, C-terminal domain | Y | N | Y | Y | Y | N |
| ENOG4103523 | polyadenylate-binding protein | Y | Y | Y | Y | Y | Y |
| ENOG410356S | Cathepsin | Y | N | Y | Y | Y | Y |
| ENOG4103598 | 40S ribosomal protein | Y | Y | Y | Y | Y | Y |
| ENOG410359F | This protein promotes the GTP-dependent binding of aminoacyl-tRNA to the A-site of ribosomes during protein biosynthesis (By similarity) | Y | Y | Y | Y | Y | Y |
| ENOG41035A9 | Dehydrogenase reductase SDR family member | Y | N | Y | N | N | Y |
| ENOG41035BZ | Produces ATP from ADP in the presence of a proton gradient across the membrane (By similarity) | Y | Y | Y | Y | Y | Y |
| ENOG41035GM | Larval storage protein (LSP) which may serve as a store of amino acids for synthesis of adult proteins | Y | Y | N | Y | N | Y |
| ENOG41035IX | Tubulin is the major constituent of microtubules. | Y | Y | Y | Y | Y | Y |
| ENOG41035KN | Required for the assembly and or stability of the 40S ribosomal subunit. | Y | Y | Y | Y | Y | Y |
| ENOG41035N8 | Glucosylceramidase (EC 3.2.1.45) | Y | Y | Y | N | Y | N |
| ENOG41035RN | alkaline phosphatase | Y | N | Y | N | Y | N |
| ENOG41035ZK | Glyceraldehyde-3-phosphate dehydrogenase | Y | Y | Y | Y | N | N |
| ENOG4103648 | Arginine kinase | Y | Y | Y | Y | Y | Y |
| ENOG41036EA | ribosomal protein S3 | Y | Y | Y | Y | Y | Y |
| ENOG41036JR | Ribosomal protein S16 | Y | Y | Y | Y | Y | Y |
| ENOG41036PA | ATP synthase | N | Y | Y | Y | Y | Y |
| ENOG41036ZV | Endonuclease_NS | Y | Y | Y | N | N | N |
| ENOG410371B | NA | Y | N | N | N | Y | N |
| ENOG410373X | Tetraspanin family | Y | N | Y | N | Y | N |
| ENOG410375W | SERine Proteinase Inhibitors | Y | Y | Y | Y | Y | Y |
| ENOG41037D0 | cytochrome C oxidase | Y | N | Y | Y | Y | Y |
| ENOG41037R1 | serine-type endopeptidase activity | Y | Y | Y | N | Y | N |
| ENOG410382G | Protein of unknown function (DUF3421) | Y | N | Y | N | Y | Y |
| ENOG41038D7 | bZIP transcription factor | N | N | Y | Y | Y | Y |
| ENOG41038K5 | serine protease | Y | Y | Y | N | Y | N |
| ENOG41039KJ | Haemolymph juvenile hormone binding protein (JHBP) | Y | N | Y | N | Y | N |

## 4. CONCLUSIONS

In the present work we described the *de novo* sequencing and assembly of transcriptomes of distantly-related, non-model insects of agronomical and medical importance. We used a computational approach to merge assemblies obtained with different algorithms and integrated our data with the eggNOG database annotations, which helped us annotate and analyze the evolution of several interesting gene families. We believe that the results provided here are not only important to shed light on fundamental biochemical aspects underlying the digestive process of the species in question, but also contribute with a large dataset that can be further used by the scientific community to design novel experiments to study genes of interest in various fields.

## ACKNOWLEDGEMENTS

This work was supported by Fundação Carlos Chagas Filho de Amparo à Pesquisa do Estado do Rio de Janeiro (FAPERJ), Fundação de Amparo à Pesquisa do Estado de São Paulo (FAPESP), Conselho Nacional de Desenvolvimento Científico e Tecnológico (CNPq) and Coordenação de Aperfeiçoamento de Pessoal de Nível Superior (CAPES).

